# Adaptation of *Drosophila* larva foraging in response to changes in food distribution

**DOI:** 10.1101/2021.06.21.449222

**Authors:** Marina E. Wosniack, Nan Hu, Julijana Gjorgjieva, Jimena Berni

## Abstract

When foraging, animals combine internal cues and sensory input from their environment to guide sequences of behavioral actions. *Drosophila* larva executes crawls, turns, and pauses to explore the substrate and find food sources. This exploration has to be flexible in the face of changes in the quality of food so that larvae feed in patches with favorable food and look for another source when the current location does not fulfill their nutritional needs. But which behavioral elements adapt, and what triggers those changes remain elusive. Using experiments and modeling, we investigate the foraging behavior of larvae in homogeneous environments with different food types and in environments where the food sources are patchy. Our work indicates that the speed of larval crawling and frequency of pauses is modulated by the food quality. Interestingly, we found that the genetic dimorphism in the *foraging* gene influences the exploratory behavior only when larvae crawl on yeast patches. While in a homogeneous substrate larvae maintain a turning bias in a specific orientation, in a patchy substrate larvae orient themselves towards the food when the patch border is reached. Therefore, by adapting different elements in their foraging behavior, larvae either increase the time inside nutritious food patches or continue exploring the substrate in less nutritious environments.

## Introduction

All living organisms need to explore their surroundings to increase their chances of finding nutritious resources. This is a challenging task in natural environments, where food quality varies both in time (e.g., seasonal effects) and space (e.g., patchy distribution). Therefore, the exploratory behavior of animals has to be flexible and adapt to environmental challenges. From the perspective of evolutionary ecology, foraging strategies have evolved to maximize lifetime fitness under distinct constraints (Stephens and Krebs, 1987) including the concentration of food inside patches (Charnov, 1976). Accordingly, several hypotheses and models have been developed to predict the optimal foraging strategy that an animal will adopt (Stephens and Charnov, 1982; Viswanathan et al., 2011), however, the effective mechanisms behind the implementation of such an optimal behavior (e.g., cognitive capacity, memory) often remain elusive (Budaev et al., 2019).

Larvae of the fruit-fly *Drosophila melanogaster* are constantly foraging and feeding to fulfill their nutritional needs for the following non-feeding pupal stage. They explore the substrate by executing sequences of crawls, pauses, and turns (Berni, 2015) and can navigate in an environment even without brain input (Sims et al., 2019). Larvae approach (or avoid) sources of odor by triggering oriented turns during chemotaxis (Gomez-Marin et al., 2011a) and can also navigate through gradients of light (Kane et al., 2013), temperature (Lahiri et al., 2011), and mechanosensory cues (Jovanic et al., 2019). Their natural habitat is decaying vegetable matter distributed in patches (Ringo, 2018), and due to food decay and intraspecific competition larvae constantly explore for new higher quality food patches. This comes at a high energetic cost since crawling behavior is very demanding (Berrigan and Lighton, 1993; Berrigan and Pepin, 1995).

The foraging behavior of *Drosophila* both in the larval and adult stages is influenced by the *foraging (for)* gene (Sokolowski, 2001; Sokolowski et al., 1997). Larvae with the rover allele crawl significantly longer paths on a yeast paste than larvae with the sitter allele, and a proportion of 70% rovers and 30% sitters is observed in natural populations (Sokolowski, 2001). Due to the higher dispersal of rover larvae, their pupae are usually found in the ground while those from sitter are usually found in the fruit (Sokolowski et al., 1986). Rover and sitter differences are also present at the adult stage: adult rover flies walk further away from a sucrose drop after feeding than sitter flies (Nagle and Bell, 1987) and are also more aggressive (Wang and Sokolowski, 2017). However, it is not known if the behavioral differences between rover and sitter larvae are present in food substrates of different compositions, nor how they behave in a patchy environment of regions with and without food (even though it has been hypothesized that rover larvae are more likely than sitter to leave a patch of food to search for a new one (Sokolowski, 2001)).

Previous studies on larval foraging focused on the behavior in homogeneous substrates, where larvae engage in a highly exploratory movement pattern if no food is available (Berni et al., 2012; Godoy-Herrera et al., 1984; Sims et al., 2019). However, the natural habitat of larvae is very patchy and it is not clear how they select feeding *vs.* exploring when the environment has food patches separated by areas without food. Previous studies have shown that larvae are more willing to leave a patch if the protein concentration is low but tend to stay in the patch if its nutritional content is adequate (Ringo, 2018). Nevertheless, these studies lack an individualized tracking of the path executed by larvae during patchy exploration.

Here we investigate the mechanisms of foraging that adapt to changes in food distribution. To address this challenge, we investigate how i) the quality of the food and ii) its distribution, homogenous *vs.* constrained in small patches, influence larval foraging. We test the effect of the rover and sitter genetic dimorphism in the different food distributions and disentangle the role of olfaction to remain in food patches using anosmic animals. By combining a detailed analysis of individual larval trajectories from behavioral experiments and computational modeling, we characterize the elements of the navigation routine and show how they adapt to a changing environment. Our results show that larvae adapt their crawling speed, turning frequency, and fraction of pauses in the face of changes in the food substrate. Importantly, larvae also trigger oriented turns towards the food at the patch interface, increasing the time larvae exploit the food inside the patch. We found only small differences in the foraging behavior of rovers and sitters in our experiments and modeling, suggesting that the foraging adaptation in changing environments is a general property of larvae.

## Results

### *Drosophila* larvae have a handedness bias during turns and decrease crawling speed in food substrates

To study the effect of different substrates in foraging larvae, we devised a behavioral assay where larvae explore different substrates with minimal external stimuli (Fig. 1A). The three different substrates (agar, sucrose, yeast) had the same agar density but distinct nutritional quality (with yeast being the richest due to its high content of protein) (Methods). Wildtype larvae from different polymorphisms – *rovers* and *sitters* – were separately recorded because of previously reported differences in foraging behavior (Sokolowski et al., 1997). We recorded the free exploratory behavior of groups of 10 3^rd^-instar larvae in large arenas (240 x 240 mm^2^) for 50 min and then tracked each individual trajectory (Risse et al., 2013). To identify salient turning points in the trajectory and obtain the distribution of turning angles of each larva, we used the Ramer-Douglas-Peucker algorithm (Methods). Larvae efficiently explored the three different substrates (Fig. 1B and Fig. S1A) by executing sequences of crawls, turns (marked as circles in the trajectories), and pauses. Interestingly, we observed that a preferential orientation – clockwise (CW) or counter-clockwise (CCW) – is present in many trajectories, and the paths described often have circular shapes (Fig. 1B).

**Figure 1.**
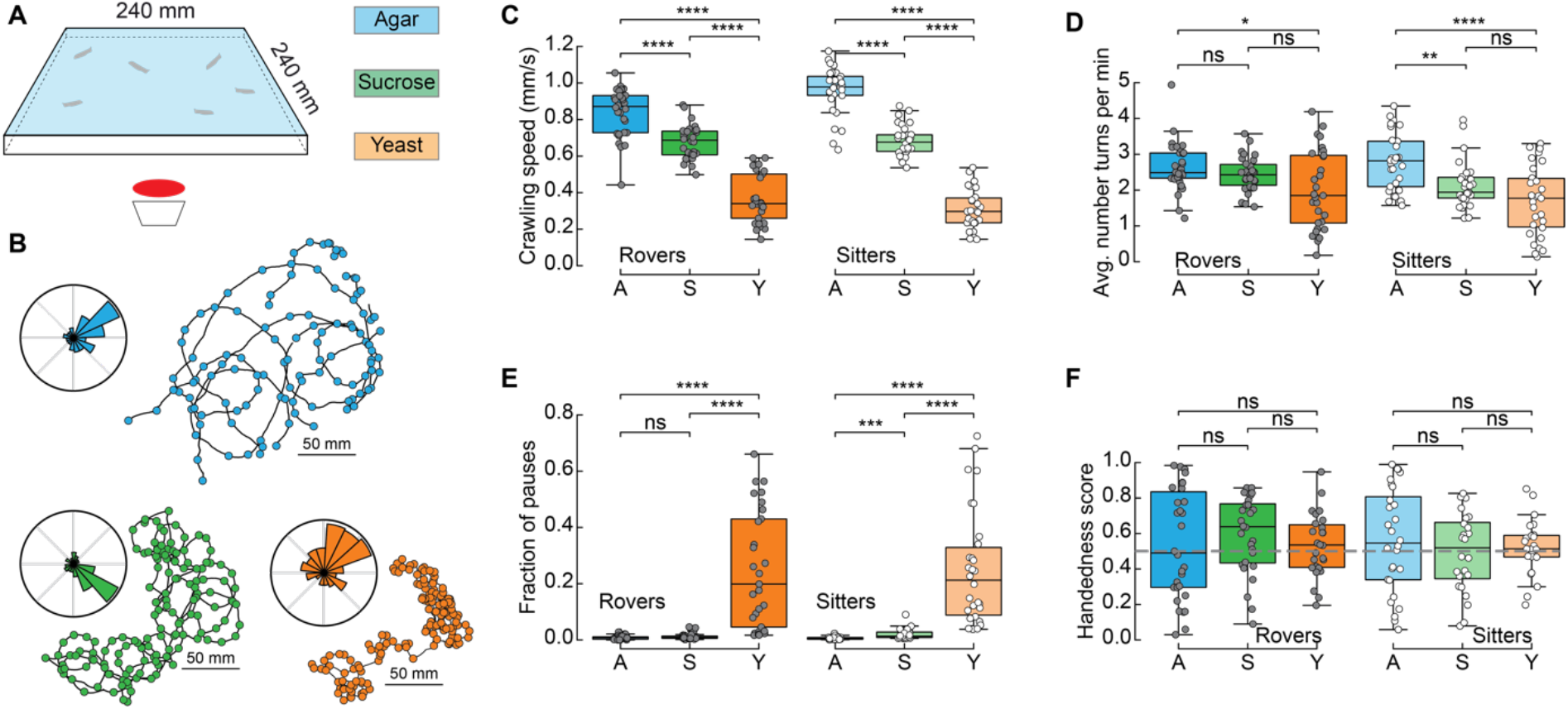
*Drosophila* larva exploratory behavior in homogeneous substrates. **A.** Experimental setup: 10 larvae of the same phenotype (rover or sitter) were placed on the top of an agar-coated arena and recorded for 50 min. Three types of substrate with the same consistency were used: agar-only (blue), sucrose (green) and yeast (orange). **B.** Sample trajectories of sitter larvae in the different substrates (top: agar, bottom left: sucrose, bottom right: yeast) with turning points identified by the RDP algorithm. Corresponding turning angle distributions are shown as an inset. **C.** Average crawling speeds of rovers (N=30,30,29) and sitters (N=29,30,30) in the different substrates: agar (A, blue), sucrose (S, green), yeast (Y, orange). Horizontal line indicates median, the box is drawn between the 25th and 75th percentile, whiskers extend above and below the box to the most extreme data points that are inside a distance to the box equal to 1.5 times the interquartile range and points indicate all data points. **D.** Average number of turns per minute registered in each trajectory. **E.** Fraction of time that larvae did not move (pauses). **F.** Handedness score. The horizontal line corresponds to a score of 0.5, i.e., an equal amount of CCW and CW turns. Welch’s t-test independent samples with Bonferroni correction. ns: 0.05 < p < 1, * 0.01 < p < 0.05, ** 0.001 < p < 0.01, *** 0.0001 < p < 0.001.

We found that the presence of food in the substrate had a strong effect on the larval crawling speed. Rover (sitter) larvae crawl on average at a speed of 0.84 mm/s, 0.68 mm/s and 0.37 mm/s (0.96 mm/s, 0.68 mm/s and 0.31 mm/s) in the agar, sucrose and yeast, respectively (Fig. 1C). In addition to changing speed, larvae suppressed turning in the food substrates, with rover (sitter) larvae executing an average of 2.65, 2.44 and 2.00 (2.79, 2.13 and 1.71) turns per minute in the agar, sucrose and yeast, respectively (Fig. 1D). Larvae also paused much more often in the yeast substrate, probably to feed (Fig. 1E).

We next quantified the individual orientation preference of each larva based on its turning angle distributions. We define the handedness score H of a larva as the number of CCW turns divided by the total number of turns in the trajectory, i.e., CCW and CW combined. Larvae with H>0.5 (H<0.5) have a bias to turn CCW (CW). Surprisingly, in both rover and sitter populations we found larvae with a very strong handedness, meaning that larvae have individual biases when turning in homogeneous environments that do not provide orientation cues in the form of sensory stimuli (Fig. 1F).

Finally, we contrasted the differences in exploratory behavior of rovers and sitters in the different homogeneous substrates (Fig. S1B-F). In particular, we were interested if sitter larvae crawled significantly less than rovers in the first 5 minutes of the recording in the food substrates, as previously observed in experiments using yeast substrates (Sokolowski, 1980). We did not find significant differences between the crawled distances of *rovers* and *sitters* in the substrates that we tested. This could be related to differences in the food preparation protocol since due to our imaging technique our substrates had to be transparent (Risse et al., 2013). Therefore, we applied a thin layer of yeast on top of the agar surface instead of thick yeast suspension as in (Sokolowski, 1980). Also, our experiments were conducted in the dark, which might influence behavior.

In summary, we have provided a detailed characterization of larval foraging behavior in homogenous substrates with different types of food. We found that the larval crawling speed and probabilities to turn and to pause are behavioral elements that are adapted according to the quality of food. Our results indicate that there are no significant differences between rover and sitter larvae in these environments.

### A phenomenological model of crawling describes larval exploratory behavior in patchy substrates

We next designed a phenomenological model to simulate larval exploratory trajectories in different substrates based on our collected data (Methods). Each type of larva (rover, sitter) has a distribution of crawling speeds *v* and probabilities to crawl P_crawl_, to turn P_turn_, and to pause P_pause_ in a given time step for each type of substrate: agar, sucrose, yeast (Fig. 2A). To capture the variability in the turning behavior, each simulated larva has its own set of parameters for the turning angle distribution according to one recorded larva. The simulated trajectories preserve the CW or CCW orientation inherited from the turning angle distributions characterized in the experiments (Fig. 2B).

**Figure 2.**
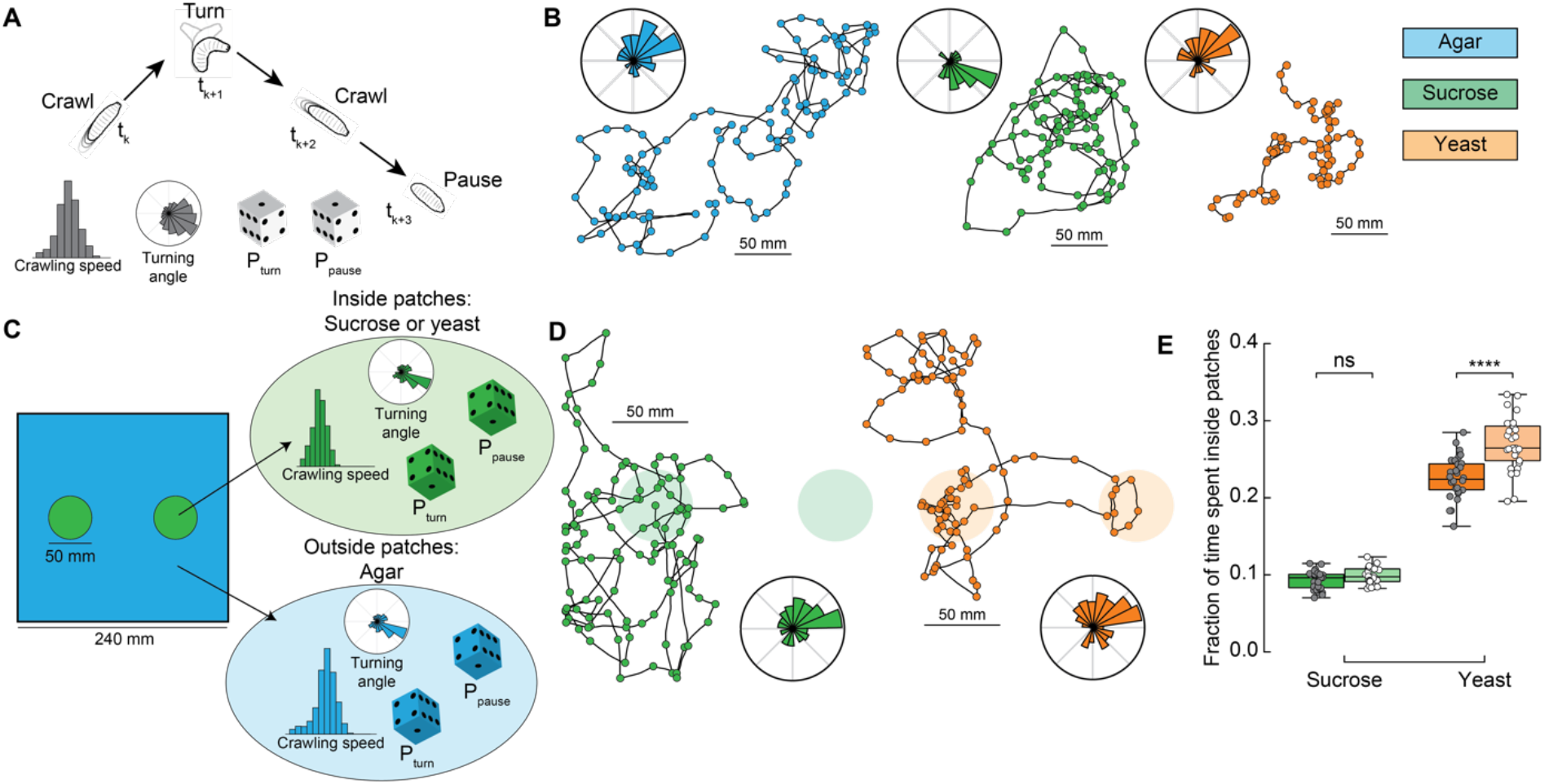
Model of larva crawling in different substrates. **A.** Simulated larva crawls at time steps tk and tk+2, turns at tk+1, and makes a pause at tk+3. Crawling speed and turning angle are sampled from normal and von Mises probability distributions, respectively. At each time step, there is a constant probability to turn P_turn_ or to pause P_pause_. **B.** Sample model trajectories and turning angle distributions of sitter larvae simulated in different homogeneous substrates: agar (left), sucrose (middle), and yeast (right). **C.** Simulations with patchy environments: food (sucrose or yeast) is distributed inside two circular regions, with agar in the remaining substrate. Crawling speeds, turning and pause probabilities are sampled from different distributions when the simulated larva is inside (green) or outside (blue) the patch. **D.** Sample model trajectories and turning angle distributions of sitter larvae in simulated patchy substrates: sucrose (left) and yeast (right) patches. **E.** Average fraction of time each simulated larva (rover, sitter, N=30) spent inside patches (sucrose, yeast) in the simulations. Horizontal line indicates median, the box is drawn between the 25th and 75th percentile, whiskers extend above and below the box to the most extreme data points that are within a distance to the box equal to 1.5 times the interquartile range and points indicate all data points. Mann-Whitney-Wilcoxon test two-sided. ns: 0.05 < p < 1, * 0.01 < p < 0.05, ** 0.001 < p < 0.01, *** 0.0001 < p < 0.001.

Using our model based on crawling behavior in homogeneous substrates, we next tested how changes in the food distribution influence the exploratory trajectories of rovers and sitters. We modeled heterogeneous environments with two circular patches of food substrate with agar substrate in the rest of the arena (Fig. 2C, see Methods). The two patches have a fixed radius (25 mm) that corresponds to the surface area of a grape (Xie et al., 2018). We simulated larval exploration of rovers and sitters in patches of two food substrates - sucrose and yeast (Fig. 2D).

We next quantified the fraction of simulation time that rovers and sitters spent inside patches. For each larva, this was averaged in 30 simulation runs (Fig. 2E and Fig. S2A, B). Inside sucrose patches, the percentage of time spent inside patches was small for both rovers and sitters (9.2% and 9.8%, respectively). These values are only slightly larger than those in a simulated environment with patches made of agar – i.e., the same speed and probabilities to turn and pause inside and outside patches – (7.47% for rovers and 6.99% for sitters – Fig. S2C, D). This result is unsurprising since in our homogeneous substrate experiments with both rover and sitter larvae had similar behavior in the agar and sucrose arenas. In simulations with yeast patches, the percentage of time spent inside patches was higher for both rovers (22.6%) and sitters (26.9%). This increase can be linked to the slower speeds and more frequent pauses in the homogeneous yeast substrate executed by rovers and sitters. In spite of non-significant differences in the crawling of rovers and sitters in the homogeneous yeast substrate (Fig. S1A-C), in our model simulated sitter larvae remained longer inside the yeast patches due to their lower (though not significantly different) average crawling speed in the homogeneous yeast substrate experiments (Table 1).

**Table 1:** Parameters of model in homogeneous and patchy substrates obtained in homogeneous substrate experiments.

| Substrate | Larva | Mean v<br>(mm/s) | Std v<br>(mm/s) | Pturn/sec | Ppause/sec |
| --- | --- | --- | --- | --- | --- |
| Agar | Rover | 0.84 | 0.13 | 0.044 | 0.0083 |
| Agar | Sitter | 0.96 | 0.13 | 0.046 | 0.0063 |
| Sucrose | Rover | 0.68 | 0.092 | 0.041 | 0.012 |
| Sucrose | Sitter | 0.68 | 0.085 | 0.035 | 0.021 |
| Yeast | Rover | 0.37 | 0.13 | 0.033 | 0.25 |
| Yeast | Sitter | 0.31 | 0.11 | 0.028 | 0.25 |

Thus far, our model does not integrate other possible mechanisms that a larva might employ to remain inside a food patch besides decreasing its crawling speed and increasing the fraction of pause events. To understand what is the effect of a food-no food interface and test the predictions of our model, we performed experiments with patchy substrates.

### At the food-no food interface, larvae turn towards the patch center

We next recorded the larval behavior in arenas with patchy substrates. We used the same size and distribution of food patches as in our simulations (Fig. 3A). Food (sucrose, yeast or an apple juice solution) was distributed inside, while agar outside patches (Fig. 3A and Methods). The concentration of the sucrose and yeast patches was the same as in the homogeneous substrate. Groups of 5 larvae of the same type (rovers or sitters) were placed inside each patch (total of 2) at the beginning of the recordings (total of 10 larvae of the same type per experiment).

**Figure 3.**
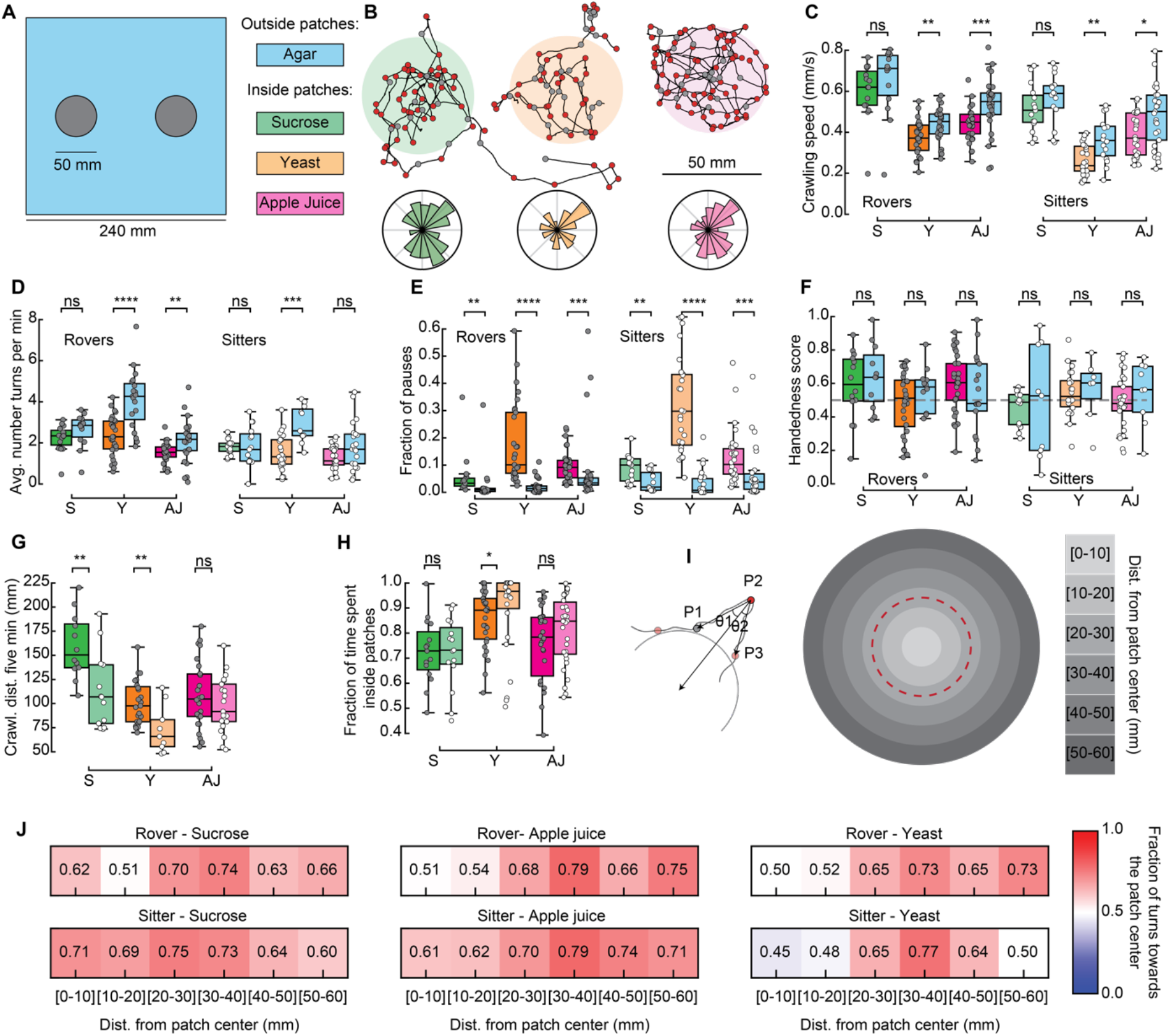
Larval exploratory behavior in patchy substrates. **A.** Experimental setup: 5 larvae of the same phenotype were placed on top of each food patch (2 patches, total: 10 larvae per experiment). Three types of food patches were tested: sucrose (green), yeast (orange) and apple juice solution (magenta). Agar was uniformly spread in the arena outside the food patches. **B.** Sample trajectories of sitter larvae in the three patch substrates with inward (outward) turns marked in red (gray) circles. Distribution of turning directions is shown on the bottom of each trajectory. **C.** Larval crawling speeds of rovers and sitters measured inside (colored bars) and outside (blue bars) food patches: sucrose (S, green), yeast (Y, orange) and apple juice (AJ, magenta). Horizontal line indicates median, the box is drawn between the 25th and 75th percentile, whiskers extend above and below the box to the most extreme data points that are within a distance to the box equal to 1.5 times the interquartile range and points indicate all data points. **D.** Average number of turns executed per minute. **E.** Fraction of time that larvae did not move (pauses). **F.** Handedness score. **G.** Total distance crawled by rover (darker colors) and sitter (lighter colors) larvae in the first 5 minutes of the recording. **H.** Fraction of time spent inside patches of rovers and sitters. **I.** Left: Identification of turning angle as inwards (θ_2_ < θ_1_, red) or outwards (θ_2_ > θ_1_, gray). Right: Circular regions with fixed distances relative to the patch center. The red dashed line represents the patch border. **J.** Relative fraction of inward turns calculated as a function of the distance from the patch center. Left: Sucrose, middle: apple juice, right: yeast patches. Top: rovers, bottom: sitters. Mann-Whitney-Wilcoxon test two-sided. Ns: 0.05 < p < 1, * 0.01 < p < 0.05, ** 0.001 < p < 0.01, *** 0.0001 < p < 0.001.

We tracked the trajectories with the same methods used in the homogeneous environment (Fig. 3B and Fig. S3A). Then, we performed the analysis separately for the two different regions: inside and outside the patches, and quantified features of the larval exploratory behavior. Inside yeast and apple juice patches, larvae crawl significantly slower than outside them (Fig. 3C). In yeast patches, both rovers and sitters execute fewer turns inside than outside (Fig. 3D). All larvae execute significantly more pauses inside the food patches than outside (Fig. 3E). We also observed that the handedness score of the larvae is less broad than in the homogeneous substrates (Fig. 3F), which may be caused by reorientations that are triggered to prevent the larva from exiting the food patch. As expected from the phenotype, sitter larvae crawled a shorter distance in the first 5 minutes of the recording in the yeast but also the sucrose substrates (Fig. 3G). In general, sitter larvae had smaller crawling speeds and executed fewer turns in the patchy environments than rovers (Fig. S3A, B). We also noticed that sitters paused more inside patches than rovers (Fig. S3C).

Our model predicted that the fraction of time spent inside patches should vary according to the substrate: larvae should remain longer inside yeast patches than inside sucrose patches (Fig. 2E). In particular, simulated sitter larvae stayed longer than simulated rovers inside yeast patches. In the experiments, the same trend was observed: for both rovers and sitters the fraction of time spent inside patches was higher in the yeast compared to both sucrose and apple juice patches (Fig. 3H). Sitter larvae had significantly higher values inside yeast patches than rovers (Fig. 3H). Nevertheless, the percentage of time the larvae spent inside patches in the experiments was very different from our model predictions. Rover (sitter) larvae remained on average 72.6% (72.3%) of the experiment inside sucrose, 85.7% (90.0%) inside yeast and 75.6% (81.3%) inside apple juice patches. Those values were much higher in the experiments than what we predicted with our simulations, and suggest that larvae might employ other mechanisms in addition to slower crawling and more frequent pauses to remain inside the food.

To gain more insight into the strategies used by larvae to remain inside the patch, we studied the distribution of turns in the food-no food interface. First, we labeled each turn as inwards or outwards depending on whether they were oriented towards or away from the patch center (Tao et al., 2020) (Fig. 3I, left). We observed that in the trajectories inward turns are consistently executed in the border of the patch for the three substrates (Fig. 3B, inward turns are shown in red). Then, we studied the fraction of turns towards the patch center as a function of the distance to the patch center (Fig. 3I, right). For the three types of substrates, the bias to turn inwards is clearly manifested when the larvae experience the patch border (patch radius: 25 mm, distance bin: 20-30 mm) (Fig. 3J). Surprisingly, the bias persists when the larva exits the patch (distance bins: 30-40, 40-50 mm). We did not consider further distance bins in our analysis because most larvae did not reach those locations in our experiments.

Therefore, we find that our model predictions are not well supported by experiments with patchy substrates. In particular, we conclude that larvae tend to orient towards the patch center when they reach the food-no food interface and this might be an important mechanism they use to remain inside the food.

### Anosmic larvae also select turns towards the patch center when reaching the food-no food border, but not on the yeast

It is well known that *Drosophila* larvae can efficiently navigate towards an odor source using chemotaxis (Gomez-Marin et al., 2011b; Schulze et al., 2015). Therefore, we wondered how much of the tendency to turn towards the patch center could be attributed to chemotaxis. Thus, we repeated the patchy experiments with mutant anosmic larvae, which have their olfactory system silenced (Orco null background), to test if they show the same distant-dependent bias when exploring the patchy substrate.

Anosmic larvae extensively explore the patchy substrate (Fig. 4A). In general, they exhibit a smaller difference in crawling speeds when comparing their behavior inside *vs*. outside of food patches (Fig. 4B). Curiously, this difference in speeds is non-significant inside *vs.* outside yeast patches. We also found that the fraction of pauses of anosmic larvae in yeast patches is smaller than that of rovers and sitters (Fig. 3G and Fig. 4D). This suggests that yeast patches do not seem to be attractive to anosmic larvae, in agreement with the lower fraction of time spent inside yeast patches relative to sucrose and apple juice patches (Fig. 4F).

**Figure 4.**
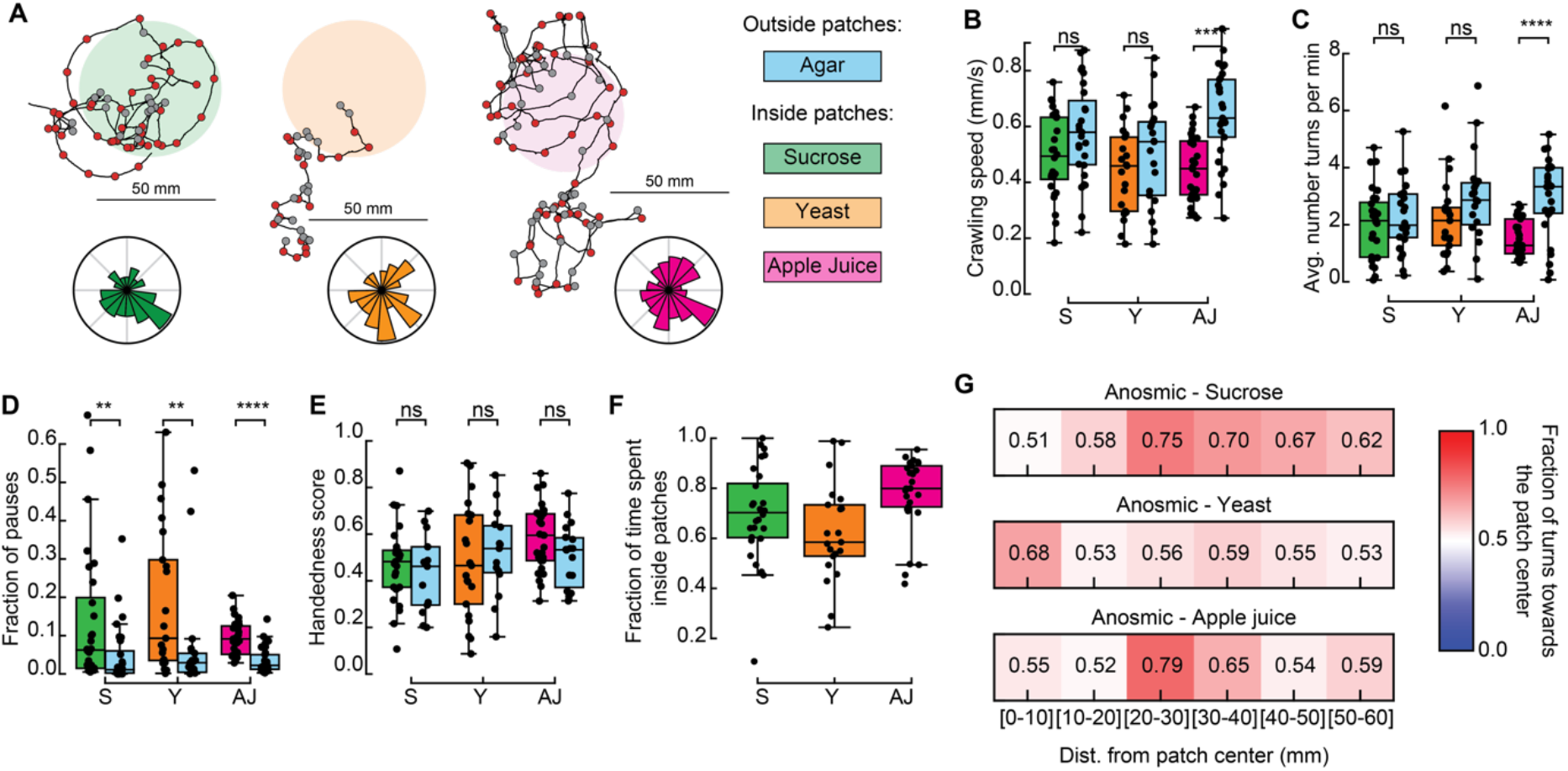
Exploratory behavior of anosmic larvae in patchy environments. **A.** Sample trajectories of anosmic larvae in the three patch substrates with inward (outward) turns marked in red (gray) circles. Distribution of turning directions is shown on the bottom of each trajectory. **B.** Crawling speeds of anosmic larvae measured inside (colorful bars) and outside (blue bars) food patches: sucrose (S, green), yeast (Y, orange) and apple juice (AJ, magenta). Horizontal line indicates median, the box is drawn between the 25th and 75th percentile, whiskers extend above and below the box to the most extreme data points that are within a distance to the box equal to 1.5 times the interquartile range and points indicate all data points. **C.** Average number of turns per minute inside and outside patches. **D.** Fraction of pauses inside and outside patches. **E.** Handedness score of anosmic larvae inside and outside patches. **F.** Fraction of time spent inside patches for different types of food. **G.** Relative fraction of inward turns calculated as a function of the distance from the patch center, top: sucrose, middle: yeast, bottom: apple juice.

Next, we investigated if anosmic larvae can bias their turns at the patch border interface without navigating odorant cues. Turns in the trajectory were labeled as inwards or outwards (as in Fig. 3I) and the fraction of turns towards the patch center was analyzed as a function of the distance away from the patch center. In sucrose and apple juice substrates, anosmic larvae consistently increased the fraction of inward turns near the patch border (Fig. 4G). This is not the case in the yeast patches, where no strong bias was detected at the patch border.

In sum, we found that anosmic larvae trigger turns towards the patch center at the food-no food interface, suggesting that chemotaxis is not the key mechanism responsible for the turning bias that increases the fraction of time larvae spend inside patches. In addition, we also found that anosmic larvae are not attracted to yeast, in contrast to sucrose and apple juice.

### Larvae employ other strategies than turning bias to remain inside a food patch

To understand the impact of the turning bias on the percentage of time that larvae spend inside patches, we included in our model (Fig. 2) a distance-dependent probability of turning towards the patch center (Fig. 5A). After drawing a turning angle from the probability distribution, the turn was implemented towards the patch center with probability *P_bias_* that depends on the distance between the current position and the center of the closest patch (Fig. 5B). For each simulated substrate, larva type, and relative distance, *P_bias_* corresponds to the fraction of turns towards the patch center quantified in our experiments (Figs. 3J and 4G).

**Figure 5.**
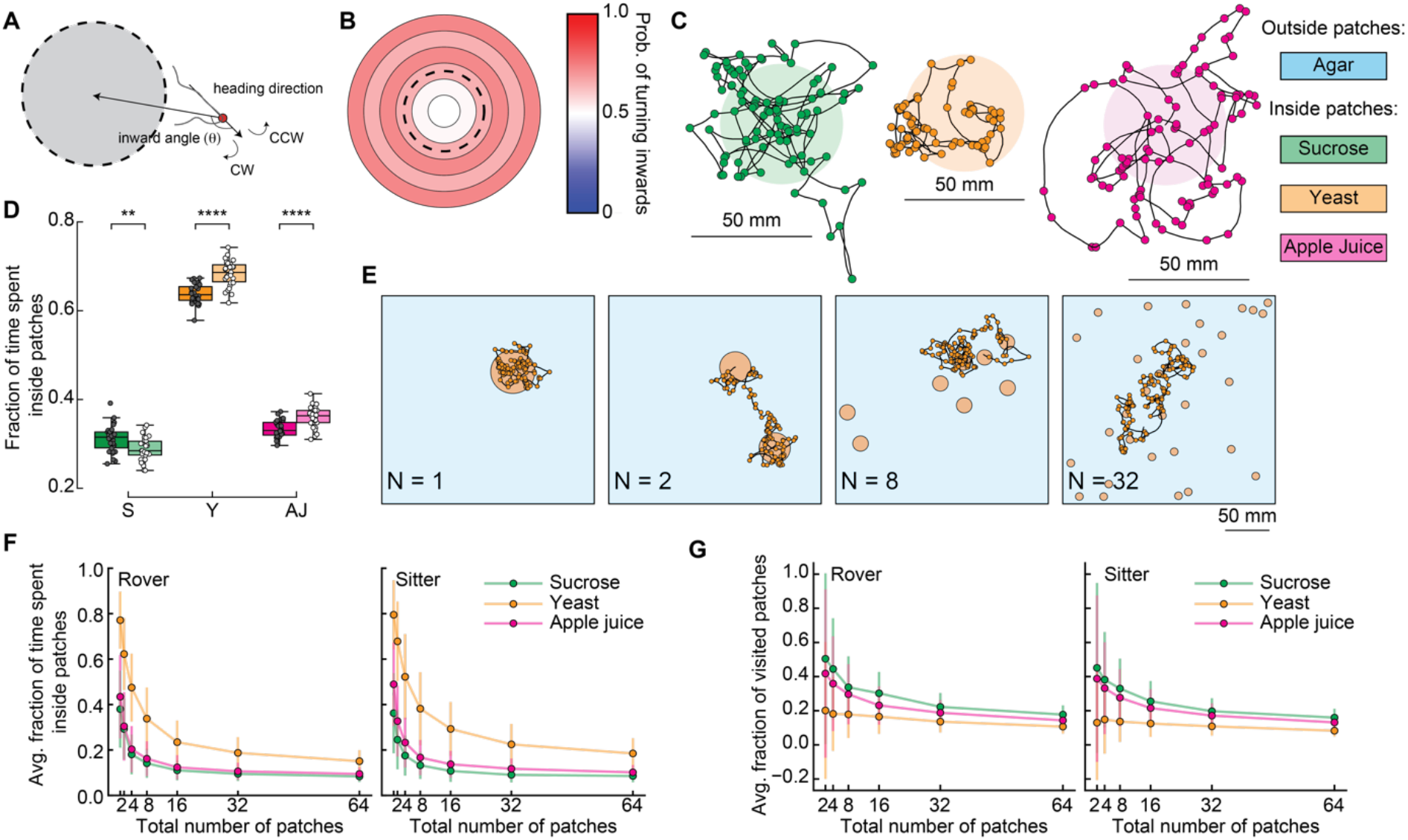
Including turns biased towards patch center in the model. **A.** Schematic showing inward turn (CW) being selected by the simulated larva. By selecting inward turns, the trajectory approaches the patch center. **B.** Spatial-dependent probability of turning towards the patch center. Each region is a concentric circle with a fixed probability of drawing inward turns (see Figure 3I, right). The dashed line shows the patch border. **C.** Sample simulated trajectories for a sitter larva with biased inward turns: sucrose patch (green), yeast patch (orange), apple juice patch (magenta). **D.** Fraction of time spent inside patches of rovers (darker colors) and sitters (lighter colors) in the different substrates: sucrose (S, green), yeast (Y, orange), and apple juice (AJ, magenta). Each point is 30 simulation runs of one larva (total: 30 larvae simulated per substrate). Horizontal line indicates median, the box is drawn between the 25th and 75th percentile, whiskers extend above and below the box to the most extreme data points that are within a distance to the box equal to 1.5 times the interquartile range and points indicate all data points. **E.** Sample trajectories of sitter larvae in environments with varying number of randomly located patches, with a fixed total area of yeast substrate being distributed (Np=1, 2, 8, 32 from left to right). **F.** Average fraction of time spent inside patches of distinct substrates (S: sucrose, green; Y: yeast, orange, and A: apple juice, magenta) for rovers (left) and sitters (right) as a function of the number of patches. Each point is the average of 30 larvae (30 simulation runs each). Bars show the standard deviation. **G.** Same as F. but for the average fraction of visited patches. Mann-Whitney-Wilcoxon test two-sided. Ns: 0.05 < p < 1, * 0.01 < p < 0.05, ** 0.001 < p < 0.01, *** 0.0001 < p < 0.001.

We observed that the simulated trajectories with this distance-dependent turning bias resemble the experimental ones much more (Fig. 5C), with larvae often returning to a patch when leaving its border. Indeed, larvae spend three times longer inside a patch in the new simulations compared to the model without biased orientations (Fig. 5D and Fig. S4): now rover (sitter) larvae remain on average 31.1% (28.9%) of the simulation inside sucrose patches and 63.8 (68.4%) of the simulation inside the yeast patches. Simulated anosmic larvae also show a gain in the ratio of time inside patches (Fig. S5A and B). Therefore, biased orientations at the patch border are an important mechanism employed by larvae to return to a food source when they detect a change in the substrate quality. Importantly, this is achieved without olfactory orientation cues, since anosmic animals can also perform biased turns (Fig. 4G). However, the ratio of time that simulated larvae remain inside patches is still smaller than what we quantified in the experiments (Figs. 3H and 4F). We reason that other mechanisms such as short-term memory or even chemotaxis can contribute to increasing the time inside the food.

### Larvae experience a trade-off between food quality and patches variability

We next asked how a further fragmentation of the food patches affects larval exploratory behavior. To test this in our model, we fixed the total area of food *S* and varied the number of patches while choosing the center coordinates for each patch randomly (Fig. 5E). To compensate for different patch radii, we adjusted the distance-dependent probability to turn inwards of each larva (Figs. 3J and 4G, see Methods).

First, we quantified the average fraction of the time spent inside patches relative to the whole simulation for the different food substrates as a function of the number of patches (Fig. 5F and Fig. S6A,B). For both rovers and sitters, larvae spend less time inside a patch as the number of patches increases (and thus the patches radii decreases) (Fig. 5F). Larvae spend longer inside patches in more nutritious environments, e.g. yeast, irrespective of the number of patches available. Interestingly, our results show that sitter larvae consistently spend more time inside yeast patches than rovers for the different number of patches (Fig. S6D). This is not observed in the sucrose or apple juice patches. Anosmic animals also spend less time inside patches when the number of patches increases, but the dependence on the quality of food is much less pronounced (Fig. S6C).

Next, we investigated the effect of different food substrates on the number of patches larvae explore. We quantified the fraction of new patches a larva visits during the simulation (discounting the source patch, since all the simulations start with the larva inside one patch) (Fig. 5G). Rovers and sitters explored more patches in the less nutritious substrate (sucrose), with a slightly higher fraction of visited patches for rovers in the sucrose and yeast patches (Fig. S6E). Anosmic larvae showed a weaker effect of the substrate on the fraction of patches visited (Fig. 5C).

Our model results suggest that by adapting their behavior to the quality and spatial distribution of food resources, larvae can either further exploit the current food source or move on to find a more profitable patch. Therefore, we predict a tradeoff between the quality of the food and the fraction of patches visited: when exploring a substrate with low-quality (high-quality) food, more (less) patches are visited. This behavior can lead to more successful foraging since larvae can adapt the amount of exploration in the substrate to the quality of the food being consumed.

## Discussion

Foraging behavior is a complex process influenced by many internal factors (locomotion style, sensory perception, cognitive capacity, age) and external variables (spatio-temporal distribution of resources, presence of predators, social interactions with co-specifics). Here, we focused on the detailed characterization of foraging in a single model organism, the fruitfly *Drosophila* larva, using extensive experiments and modeling. This allowed us to study the role of both internal and external factors on foraging: i) genetics (*rovers, sitters*, and later *orco* null anosmic animals), ii) food quality (agar, yeast, sucrose and apple juice) and iii) food spatial distribution (homogeneous and heterogeneous environments).

We systematically investigated larval exploratory behavior first in experimental arenas with homogeneously distributed food. Larval crawling speed, turning frequency and fraction of pausing events adapted according to the quality of the food substrate (Fig. 1C-E). We observed that larval trajectories often had a circular shape, revealing an individualized preference for a given turning direction in the absence of direction cues, which we quantified as the larval handedness (Fig. 1B, F). The population variability in the handedness has been quantified in adult flies (Buchanan et al., 2015a), but to our knowledge not yet at the larval stage. In adult walking flies, individual preferences of turning left or right in maze tests have been shown to persist across days (Buchanan et al., 2015b) and recently have been linked to anatomical differences in the synaptic distribution of bottleneck neurons downstream of the central complex (Skutt-Kakaria et al., 2019). It is interesting that at the larval stage the turning bias is also present, and we speculate that the mechanisms for it could be different.

It is expected that animals change their foraging behavior depending on the quality and spatial distribution of food, with more localized exploitation of resources where they are abundant and a more exploratory behavior when resources become scarce (Humphries et al., 2010). We tested this in a phenomenological model of larval foraging behavior in patchy substrates (Humphries et al., 2010) (Fig. 2). We reasoned that crawling speed, turning frequency and fraction of pauses are the behavioral elements that adapt when the larva crosses the food-no food interface at the patch boundary. To quantify the food exploitation, we measured the fraction of the time each larva spent inside the patches. We found that decreasing the speed and turning frequency and increasing the fraction of pauses is not sufficient to explain why larvae remain inside the food for longer periods.

In experiments with patchy substrates, we found that larvae spend a longer time inside food patches than what we predicted with our model (Fig. 3H). The lack of agreement between the experiments and our model was not surprising, since the latter does not include additional mechanisms that could guide the larva back to the patch when it leaves it, such as chemotaxis (Gomez-Marin et al., 2011a). Since in chemotaxis larvae redirect their turns towards a source of odor, we classified each turn in their recorded trajectory as towards or away from the patch center. We observed that the fraction of inward turns is very high around the patch border (Fig. 3J). To test whether larvae could redirect their turns towards the food when exiting it only using chemotaxis, we repeated the experiments with anosmic mutants. Surprisingly, in sucrose and apple juice substrate anosmic larvae bias their turns towards the patch center when in the neighborhood of the patch border (Fig. 4G). Therefore, this reorientation at the border does not seem to rely only on chemotaxis.

By including a distance-dependent probability to turn inwards in our model, we observed that simulated larvae managed to remain inside the food for longer periods closer to experimental observations (Fig. 5D). Therefore, the ability to direct turns towards the food seems to be another important element in the foraging routine necessary to explore patchy environments.

Overall, we found both in our experiments and modeling that larvae spend less time exploiting patches of less nutritious food (e.g., sucrose). What could be the effect of this when several patches are available in the substrate? Our model results predict that more patches are visited by larvae when their food quality is lower (Fig. 5G). In natural environments, this enhances the chances that larvae will eventually find a better food source in the surroundings.

Interestingly, the differences we found in the foraging behavior of rovers and sitters are not as drastic as previously reported, where the length of the path of rovers was roughly twice that of sitters when crawling in a yeast paste for 5 minutes (Sokolowski, 2001). In the homogeneous agar, sucrose, and yeast substrates, we did not observe significant differences in the path length of rovers and sitters (Fig. S1). Curiously, when the food is constrained inside patches, we observed significantly shorter crawling paths of sitters in sucrose and yeast patches (Fig. 3G) and also slower crawling speeds, fewer turns per minute and more pauses (Fig. S3). It is possible that the presence of a patch border in our experiments, as well as a different preparation of the yeast substrate than the one from the original publications, can trigger the observed behavioral differences.

In summary, we have identified a set of behavioral elements — the crawling speed, frequency and biasing of turns, and fraction of pauses — that adapt when larvae explore environments with a patchy distribution of food sources. This adaptation leads to an efficient substrate exploration, as larvae either increase the time inside nutritious food patches or continue exploring the substrate depending on the local quality of food.

## Acknowledgments

MW was supported by a Capes-Humboldt postdoctoral fellowship. JB was funded by a Sir Henry Dale fellowship from the Wellcome Trust and Royal Society 105568/Z/14/Z. JG was funded by the Max Planck Society. The authors thank Dr. Bertram Gerber and his lab for helpful discussions during early stages of this work.

## Material and Methods

### 1. Animals

Rover and sitter flies were a gift of Marla Sokolowski (University of Toronto) and Orco^[2]^ from Bloomington stock center (stock 23130). Flies were allowed to lay eggs for one day in standard corn meal food, which consists of 420 g of cornmeal; 450 g of dextrose; 90 g of yeast; 42 g of agar; 140 ml of 10% Nipagin in 95% EtOH; 22 ml of propionic acid and 6.4 l of water. Larvae that were 72 hrs old were collected for the experiment.

### 2. Larva tracking

We recorded movies of larval exploratory behavior in arenas with minimal external stimuli – the recordings were made in the dark with a constant temperature of 25 °C. Each trial lasted 50 min and the larvae were simultaneously tracked in a 240 x 240 mm^2^ arena with a 2 mm thick layer of 0.4% agar-based coating (see the protocol of substrate preparation below).

At each trial, 10 young third-instar larvae (72 – 80 h) of approximately the same size were washed to remove traces of food and allowed to crawl freely for 5 min on a clean 0.4% agar coated plate before being transferred to the arena. We used a Frustrated Total Internal Reflection (FTIR)-based imaging method to record the larval exploratory behavior (Risse et al., 2013). Movies (duration 50 min) were recorded at 2 fps with a Basler acA2040-180km CMOS camera using Pylon and StreamPix software, mounted with a 16 mm KOWA IJM3sHC.SW VIS-NIR Lens and 825 nm high performance longpass filter (Schneider, IF-093). The resolution was 2048 x 2048 px^2^ to obtain forward movement displacements and actual pause-turns that are recorded accurately rather than to include ‘flickering’ movements associated with peristaltic movements.

**Table:** number of larvae per recording.

| Substrate | Number of trials | Average number of larvae per trial | Total larvae |
| --- | --- | --- | --- |
| Agar homogeneous | 3 rovers | 10 rovers | 30 rovers |
|  | 3 sitters | 10 sitters | 29 sitters |
| Sucrose homogeneous | 3 rovers | 10 rovers | 30 rovers |
|  | 3 sitters | 10 sitters | 30 sitters |
| Yeast homogeneous | 3 rovers | 10 rovers | 29 rovers |
|  | 3 sitters | 10 sitters | 30 sitters |
| Sucrose patches | 2 rovers | 8 rovers | 15 rovers |
|  | 2 sitters | 8 sitters | 15 sitters |
|  | 3 anosmic | 10 anosmic | 28 anosmic |
| Yeast patches | 3 rovers | 10 rovers | 30 rovers |
|  | 3 sitters | 10 sitters | 27 sitters |
|  | 3 anosmic | 10 anosmic | 21 anosmic |
| Apple juice patches | 3 rovers | 10 rovers | 30 rovers |
|  | 3 sitters | 10 sitters | 30 sitters |
|  | 3 anosmic | 10 anosmic | 29 anosmic |

### 3. Substrate preparation

The following food substrates were prepared for our experiments, and stored refrigerated for up to one day:

a. Agar substrate: 0.8 g of agar was melted in 200 ml of distilled water;
b. Sucrose substrate: 0.8 g of agar with 3.42 g of sucrose was melted in 200 ml of distilled water;
c. Apple juice substrate: 0.8 g of agar with 0.342 g of sucrose and 5 ml apple juice was melted in 195 ml of distilled water;
d. Yeast substrate: 0.8 g of agar was melted in 200 ml of distilled water with a layer of 5 ml of 20% yeast in water on top.

In the case of agar and sucrose homogeneous substrates, the solution was homogeneously spread on top of the acrylic arena and we waited for it to reach room temperature before transferring the larvae to the arena. Yeast homogeneous arenas were obtained by spreading 5ml of 20% yeast in water with a soft metallic disk. For sucrose or apple juice patchy arenas, first, the agar solution was homogeneously spread in the acrylic arena. When the solution cooled down, two holes in the agar were made at fixed positions (Fig. 3A) using circular-shaped Petri dishes with a 25 mm radius. We carefully removed the agar inside the holes and transferred the food solutions to the holes with the same thickness as the agar around them. For yeast patches, 100 μl of yeast solution was placed on a 25-mm-radius metal disc and printed on the agar. This was repeated to make two equidistant patches.

### 4. Descriptive statistics of larval trajectory

The data (*x,y* coordinates of individual larvae) were extracted from the behavioral movies using the FIM track free software (Risse et al., 2017). We used a Kalman filter to the (*x,y*) coordinates of each larva (code available at github). The position of each larva in video frame *j* is represented as the vector:

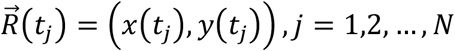

where *x*(*t_j_*) and *y*(*t_j_*) are the centroid coordinates, *t_j_* = *j*Δ*t* (*j* = 1, …, *N*), Δ*t* = 0.5 s, and *N* = 6000 is the number of frames continuously recorded during the experiment. We defined the following quantities that were used in our analysis.

Velocity:

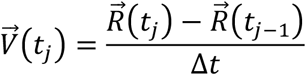

Heading:

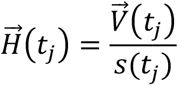

Scalar speed:

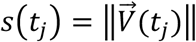

Instantaneous turn rate:

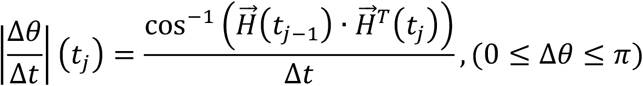

Next, the Ramer-Douglas-Peucker algorithm (https://pypi.org/project/rdp/) was used to simplify the larval trajectories and therefore identify the locations where larvae executed turns. After visual inspection of the simplified trajectories, we fixed the distance dimension *ε* = 2.5 mm to the analysis with agar, sucrose, and apple juice and *ε* = 1.25 mm to the yeast analysis.

With turning points identified in the trajectory, the turning angles were obtained in the range [−*π*, *π*] using the atan2 function in python. As a convention, clockwise turns were in the range [−*π*, 0) and counter-clockwise turns in the range [0, *π*]. The handedness index of each larva was obtained as:

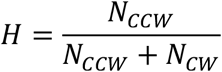

where *N_CCW_* is the number of counter-clockwise turns in the trajectory and *N_CW_* the number of clockwise turns. Thus, if *H* > 0.5 (*H* < 0.5) the larva has a bias to execute more counter-clockwise (clockwise) turns.

From the turning points identified by the RDP algorithm, we built a vector that registers 1 in the time points where turns were registered and 0 otherwise. The length of this vector is the number of frames in the recording. Next, we applied a rolling window of 120 frames (one minute) to this vector and summed the elements within the window. Then, we averaged the number of turns registered within each one-minute window to obtain the average number of turns per minute.

### 5. Patch radius and center coordinates

We used imageJ to determine the center and radius of each patch in the experiments. A frame of the recording was adjusted for contrast and brightness until the borders of the patch became visible. Circular regions of interest were drawn for each patch and the center coordinates and radius were obtained.

### 6. Classification of turns as towards the patch center

Let 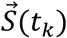 be the trajectory simplified by the RDP algorithm, where each point is a turning point of the original trajectory. To classify the *k*th turn in the trajectory as inwards or outwards, we define the following vectors:

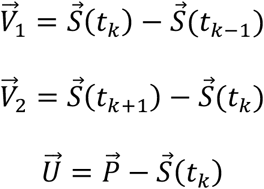

where 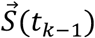 and 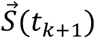 are the previous and the following turning locations and 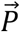 is the center of the patch that is closest to 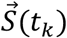. The following angles are then computed (Fig. 3I left):

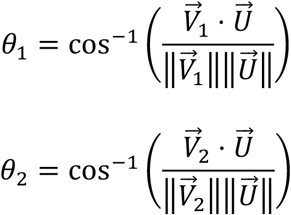

and the turn at 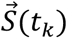 is classified as inwards (outwards) if *θ*_2_ < *θ*_1_ (*θ*_2_ > *θ*_1_) (Tao et al., 2020).

### 7. Model

#### i) Homogeneous Substrate

The simulated crawling substrate has rigid boundaries and the same dimensions as the behavioral arenas used in the experiments (240 x 240 mm^2^). At each time step *t_k_* the simulated larva can be at one of three different states (Fig. 2A):

1. with probability P_crawl_ crawling with speed *v*(*t_k_*) > 0 sampled from a normal distribution;
2. with probability P_turn_ turning an angle *θ*(*t_k_*) sampled from a von Mises distribution;
3. with probability P_pause_ paused (*v*(*t_k_*) = 0).

The parameter values and distributions were obtained from our experimental data of larval crawling in homogeneous substrates and are unique for each type of larva and substrate (Table 1). Crawl, turn or pause events are registered with a constant probability per time step (P_crawl_ = 1 – (P_turn_ + P_pause_)) and the simulation duration is the same as our behavioral recordings (50 min). To capture the variability in the turning behavior, each larva was simulated with its own set of parameters for the turning angle distribution according to one recorded larva (with an average of 30 sitter and 30 rover larvae recorded at each type of substrate). The RDP algorithm was then used to identify salient turning points in the simulated trajectory (Fig. 2B).

#### ii) Patchy substrate

##### Without biased turns towards the food

We modeled patchy environments initially with two circular patches (radius 25 mm) of food substrate (sucrose or yeast) with agar substrate in the rest of the arena (Figure 2C). Crawling speed and probabilities to turn or pause are drawn based on the current position of the simulated larva. The parameters are sampled from the corresponding food experiment when the larva is inside a patch, and sampled from the agar experiment when the larva is outside the patch. The turning angle distribution of each simulated larva corresponds to one from the recordings in the agar substrate. The same turning angle probability distribution is used whether the larva is inside or outside the patch.

##### With biased turns towards the food

Except for the choice of turning angles, the model is the same as the one described above. The biased choice of turns towards the food follows the implementation in (Tao et al., 2020). After drawing a turning angle from the von Mises probability distribution, the turn direction was chosen such that the larva points towards the patch center with probability *P_bias_* that depends on the distance between the current position relative to the center of the closest patch (Figure 5B). When the simulated larva was further than 60 mm away from the closest patch center, no bias was applied in the turning direction since the data was very sparse in this region (most larvae never crawled such long distances away from the patch of food in the experiments). Each turn is defined by a set of three points {p_1_, p_2_, p_3_} where p_1_ is where the turn initiates, p_2_ is the end location of a left turn and p_3_ the end location of a right turn. Three movement vectors that characterize the turn options (to the left or to the right) are defined as:

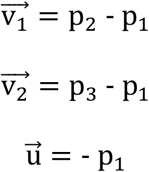

We next calculate the angle *θ* the larval trajectory makes with the inward vector 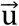 when a turn to the left (p_2_) or to the right (p_3_) is made. The inward turn is the turn that results in the smallest *θ* (as shown in Fig. 5A).

##### With more patches

We fixed the total surface area of food to be distributed in N patches as S = 2*π*R^2^, where R = 25 mm is the radius of the patches from the previous simulations and experiments. Then, the radius of each N^th^ patch is given by 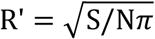. The model then is the same as in the case of two patches except that the distances in the distance-dependent probability to turn inwards are adjusted for smaller patch radius, by multiplying the distance values by R’/R.

#### iii) Model parameters

**Table 2:**
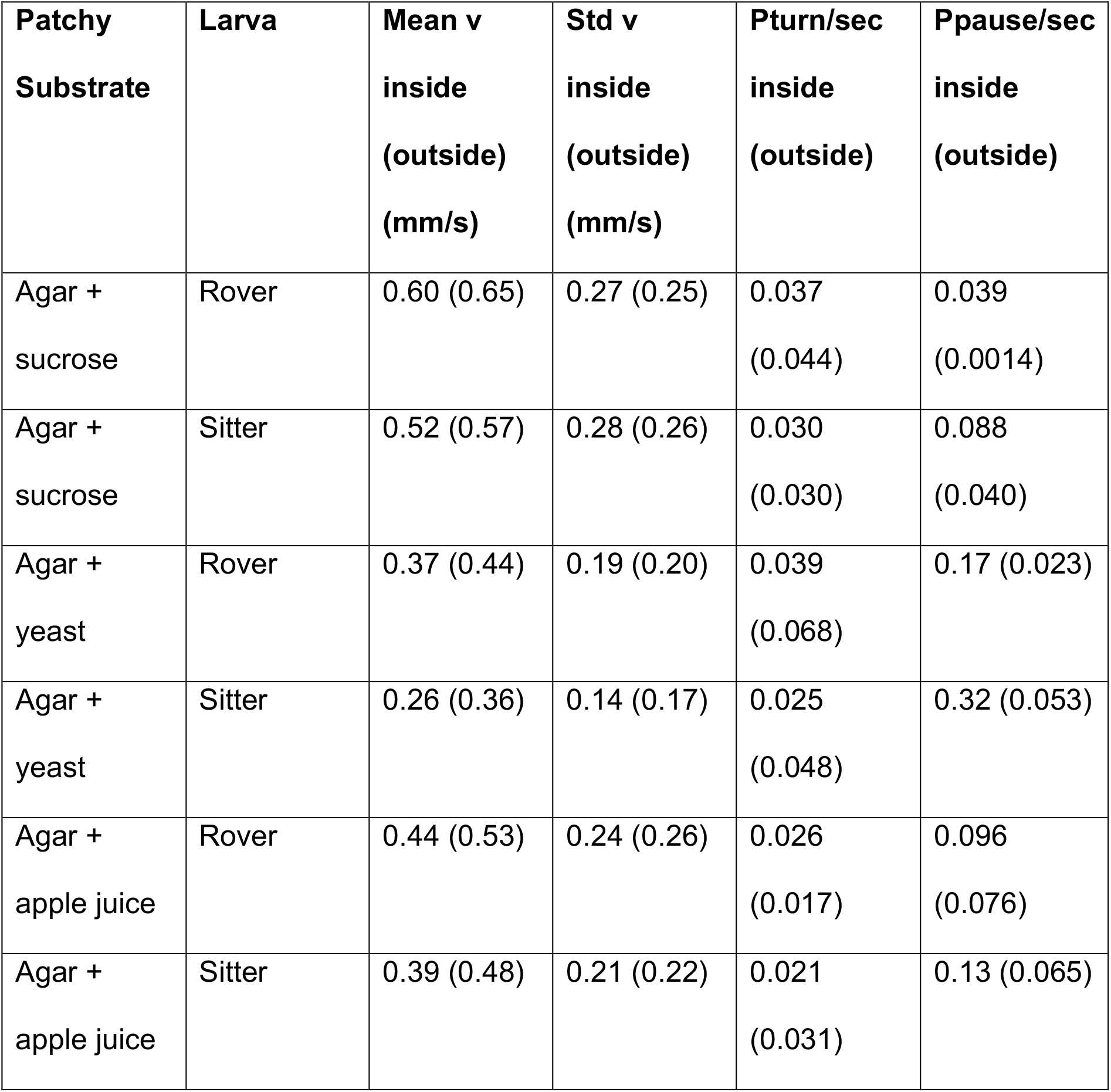
Parameters of corrected model in patchy substrates obtained in patchy substrate experiments.

| **Patchy**<br>**Substrate** | **Larva** | **Mean v**<br>**inside**<br>**(outside)**<br>**(mm/s)** | **Std v**<br>**inside**<br>**(outside)**<br>**(mm/s)** | **Pturn/sec**<br>**inside**<br>**(outside)** | **Ppause/sec**<br>**inside**<br>**(outside)** |
| --- | --- | --- | --- | --- | --- |
| Agar +<br>sucrose | Rover | 0.60 (0.65) | 0.27 (0.25) | 0.037<br>(0.044) | 0.039<br>(0.0014) |
| Agar +<br>sucrose | Sitter | 0.52 (0.57) | 0.28 (0.26) | 0.030<br>(0.030) | 0.088<br>(0.040) |
| Agar +<br>yeast | Rover | 0.37 (0.44) | 0.19 (0.20) | 0.039<br>(0.068) | 0.17 (0.023) |
| Agar +<br>yeast | Sitter | 0.26 (0.36) | 0.14 (0.17) | 0.025<br>(0.048) | 0.32 (0.053) |
| Agar +<br>apple juice | Rover | 0.44 (0.53) | 0.24 (0.26) | 0.026<br>(0.017) | 0.096<br>(0.076) |
| Agar +<br>apple juice | Sitter | 0.39 (0.48) | 0.21 (0.22) | 0.021<br>(0.031) | 0.13 (0.065) |

---

**Figure S1.**
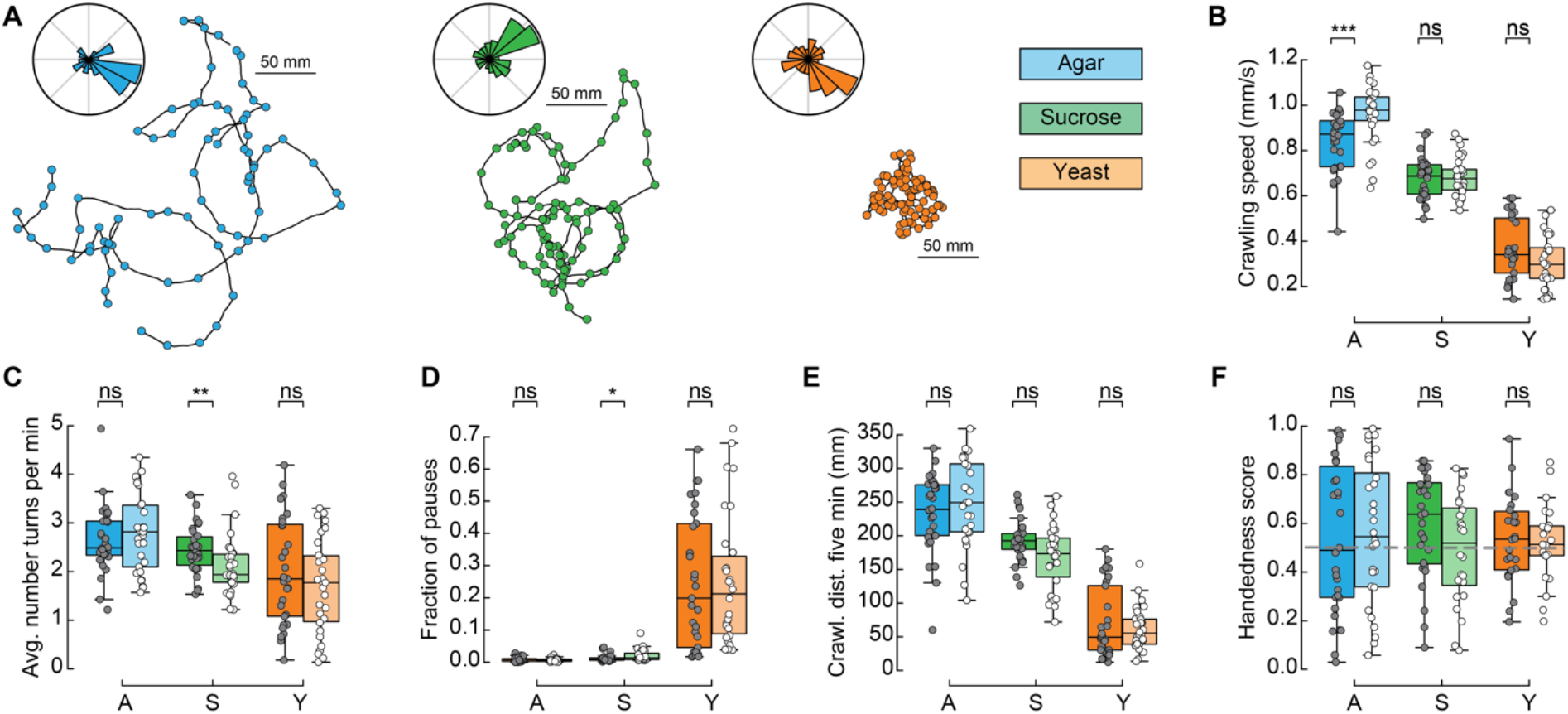
Comparison between rover and sitter behavior in different substrates. **A.** Sample trajectories of rover larvae in different substrates with the respective turning angle distributions in the inset. Left: agar, middle: sucrose, right: yeast. **B.** Crawling speeds in the agar (A, blue), sucrose (S, green) and yeast substrate (Y, orange). Darker colors are used to label rovers data, lighter colors label sitters data. Horizontal line indicates median, the box is drawn between the 25th and 75th percentile, whiskers extend above and below the box to the most extreme data points that are within a distance to the box equal to 1.5 times the interquartile range and points indicate all data points. **C.** Average number of turns executed by the larvae per minute. **D.** Fraction of pauses in the recording. **E.** Crawled distance in the first 5 minutes of the recording. **F.** Handedness score. Mann-Whitney-Wilcoxon test two-sided. ns: 0.05 < p < 1, * 0.01 < p < 0.05, ** 0.001 < p < 0.01, *** 0.0001 < p < 0.001.

**Figure S2.**
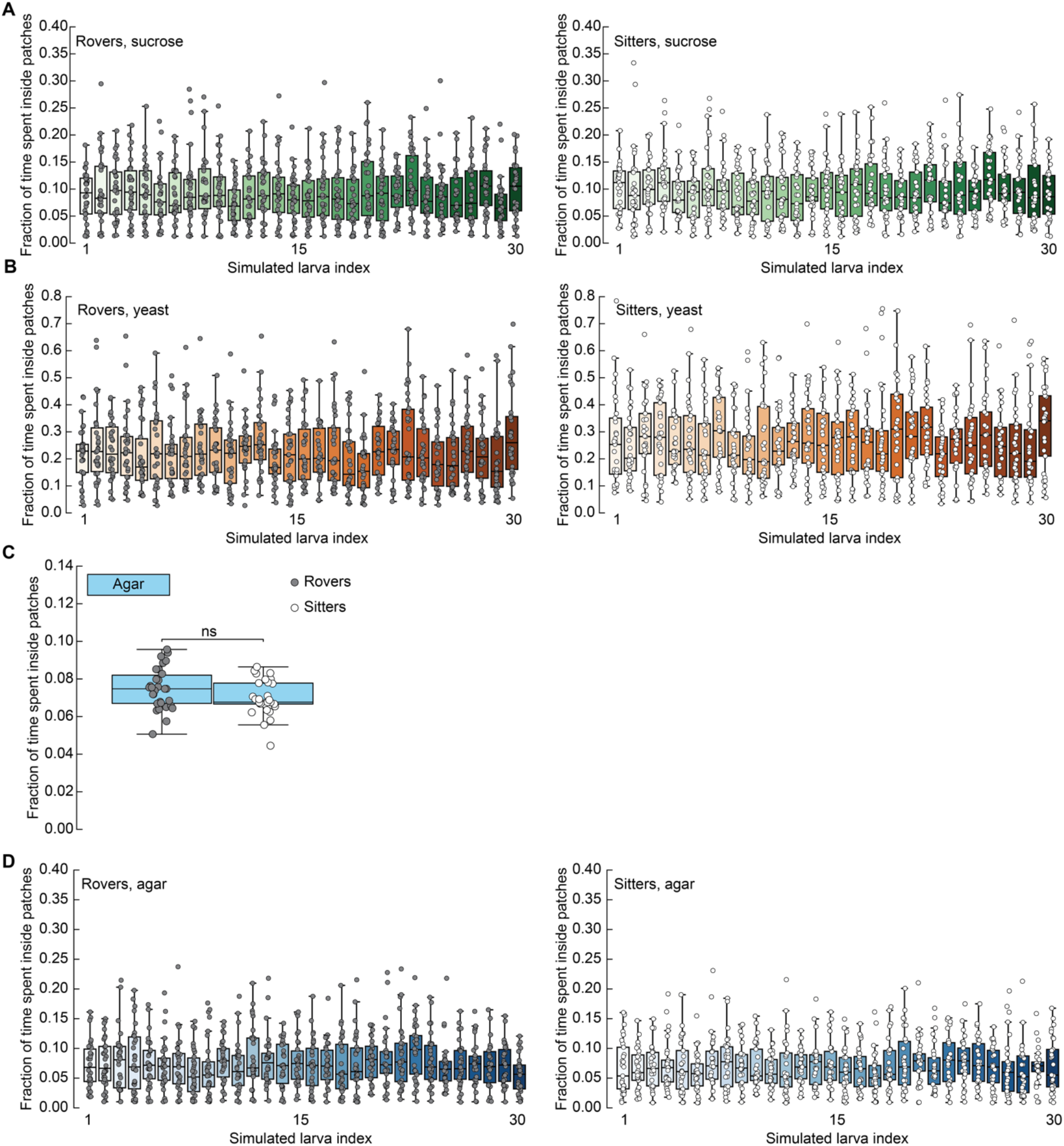
Fraction of time spent inside patches. **A.** Each simulated rover (left) and sitter (right) larva (bars with different color shades, N=30) has a fixed turning angle distribution with parameters corresponding to one rover/sitter from the agar experiments. N=30 simulation runs of each larva were performed in the same environment with sucrose (green), yeast (orange), and apple juice (purple) patches. Horizontal line indicates median, the box is drawn between the 25th and 75th percentile, whiskers extend above and below the box to the most extreme data points that are within a distance to the box equal to 1.5 times the interquartile range and points indicate all data points. **B.** Same as A. but for yeast patches. **C.** Average fraction of time spent inside patches of rovers and sitters in agar patches, where the same parameters are used for inside and outside the patches. **D.** Same as A. for agar patches.

**Figure S3.**
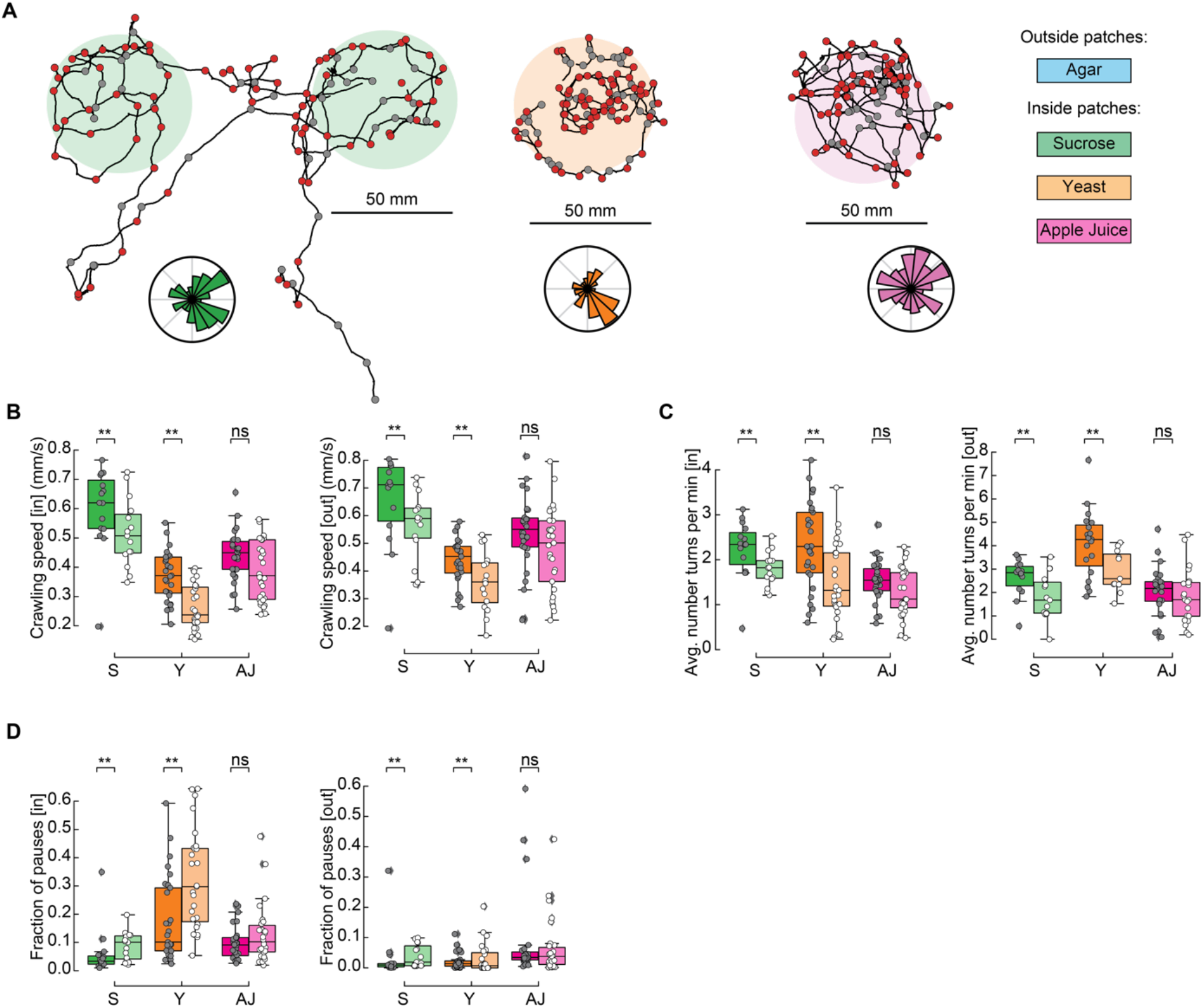
Comparison between rover and sitter behavior in patchy substrates. **A.** Sample trajectories of rover larvae in the three patch substrates: sucrose (green), yeast (orange) and apple juice (magenta). Inward (outward) turns are marked in red (gray) circles. Distribution of turning directions is shown on the bottom of each trajectory. **B.** Left: crawling speed inside patches: sucrose (S, green), yeast (Y, orange), apple juice (AJ, magenta). Data from rover (sitter) larvae shown in darker (lighter) colors. Right: crawling speed outside patches. Horizontal line indicates median, the box is drawn between the 25th and 75th percentile, whiskers extend above and below the box to the most extreme data points that are within a distance to the box equal to 1.5 times the interquartile range and points indicate all data points. **C.** Average number of turns per minute inside (left) and outside (right) patches. **D.** Fraction of pauses inside (left) and outside (right) patches.

**Figure S4.**
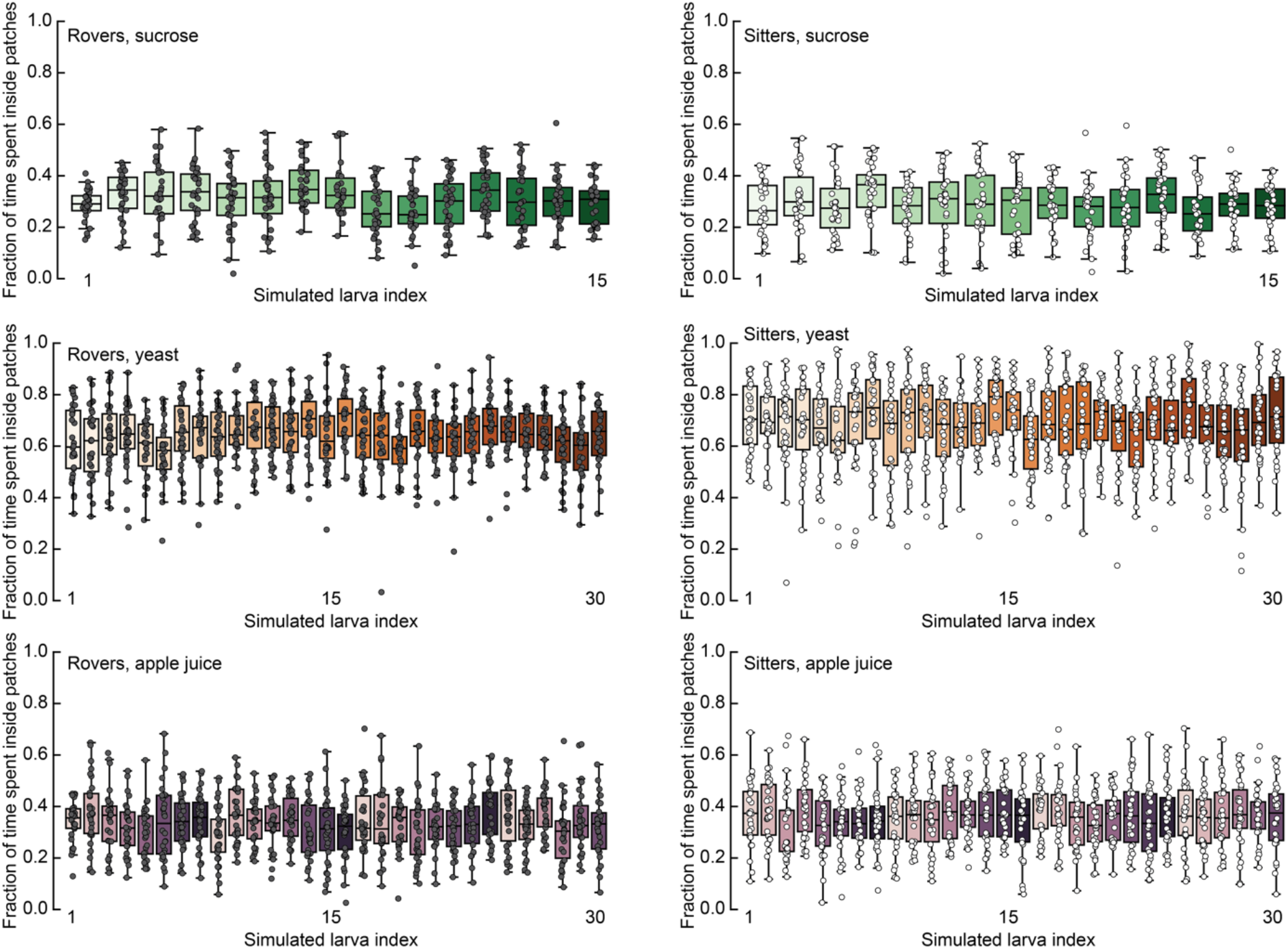
Fraction of time spent inside patches for the individual larvae. Each rover/sitter larva (bars, N=15 for sucrose, N=30 for yeast and apple juice) has a fixed turning angle distribution with parameters corresponding to one rover/sitter from the agar experiments. N=30 simulation runs were performed in the same environment with sucrose (green), yeast (orange), and apple juice (purple) patches. Horizontal line indicates median, the box is drawn between the 25th and 75th percentile, whiskers extend above and below the box to the most extreme data points that are within a distance to the box equal to 1.5 times the interquartile range and points indicate all data points.

**Figure S5.**
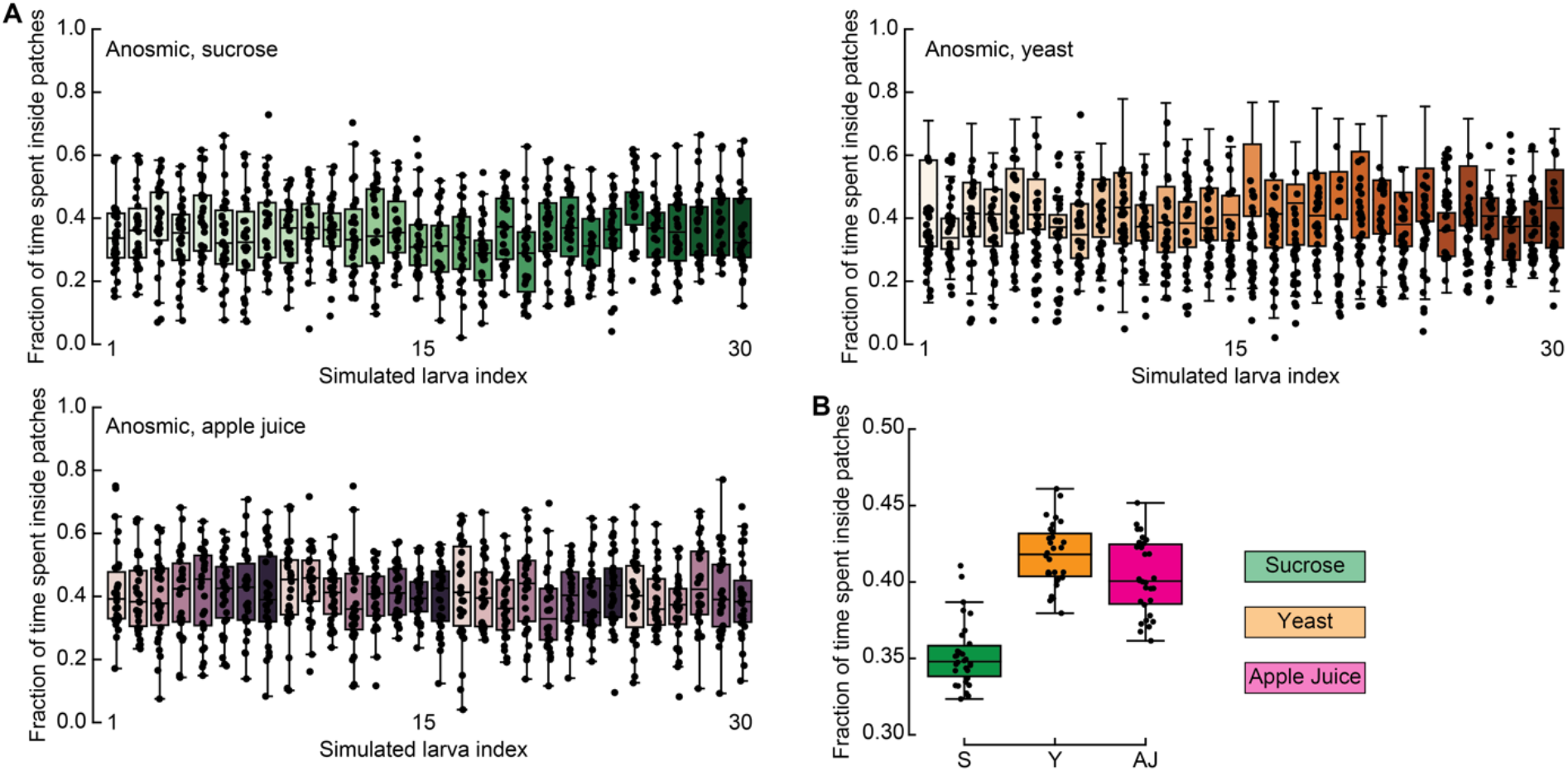
Fraction of time spent inside patches for the individual anosmic larvae. **A.** Each anosmic larva (bars, N=30 for sucrose, yeast and apple juice) has a fixed turning angle distribution with parameters corresponding to one anosmic larva from the agar experiments. N=30 simulation runs were performed in the same environment with sucrose (green), yeast (orange), and apple juice (purple) patches. Horizontal line indicates median, the box is drawn between the 25th and 75th percentile, whiskers extend above and below the box to the most extreme data points that are within a distance to the box equal to 1.5 times the interquartile range and points indicate all data points. **B.** Average fraction of time spent inside patches of anosmic larvae in sucrose (S, green), yeast (Y, orange), and apple juice (AJ, magenta) patches. Mann-Whitney-Wilcoxon test two-sided. Ns: 0.05 < p < 1, * 0.01 < p < 0.05, ** 0.001 < p < 0.01, *** 0.0001 < p < 0.001.

**Figure S6.**
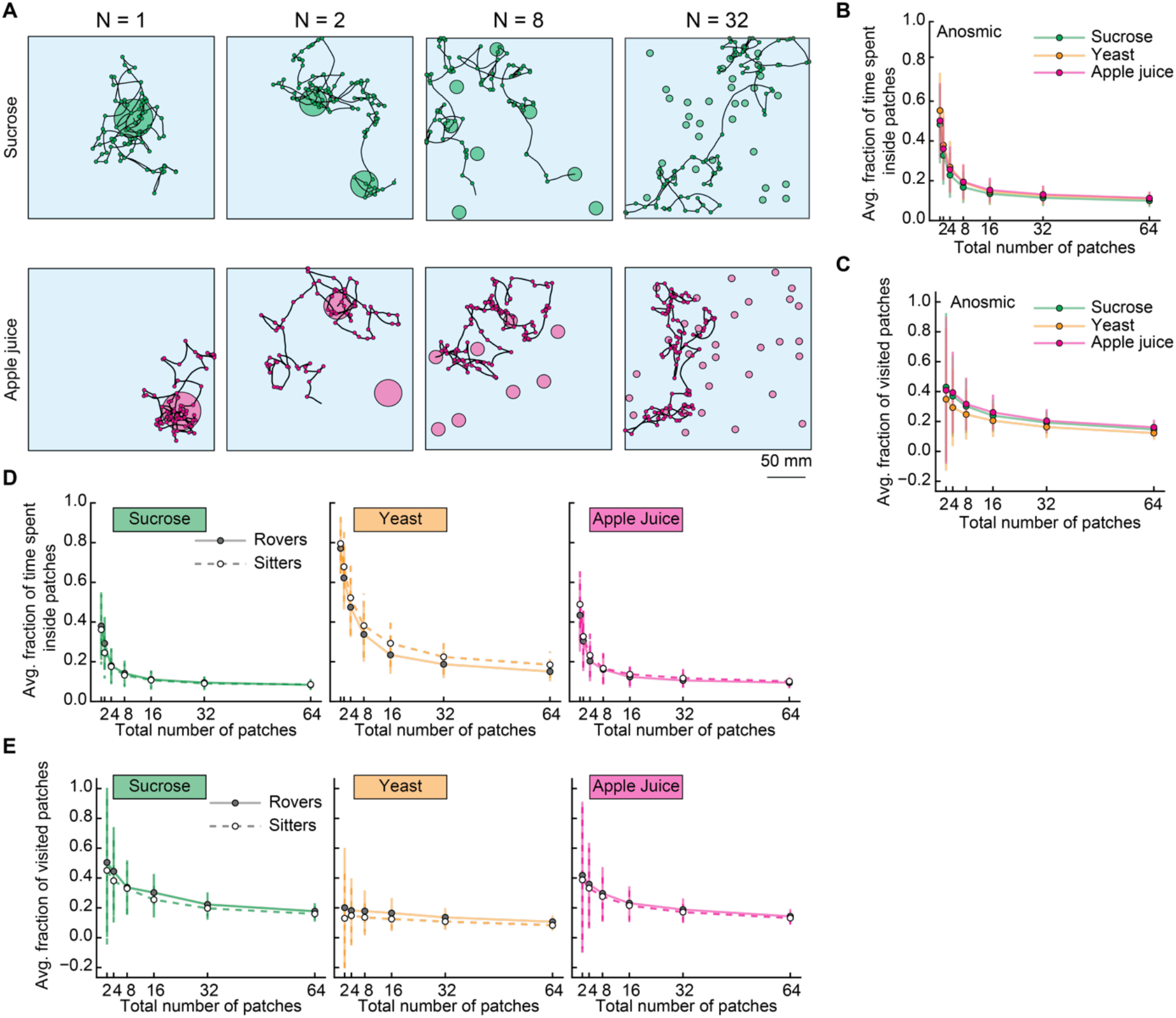
Simulations with varying number of patches. **A.** Sample trajectories of sitter larvae in sucrose (top) and apple juice (bottom) patchy arenas with varying number of patches (Np=1, 2, 8, 32 from left to right). **B.** Average fraction of time spent inside patches of distinct substrates (S: sucrose, green; Y: yeast, orange, and A: apple juice, magenta) for anosmic larvae as a function of the number of patches. Each point is the average of 30 larvae (30 simulation runs each). Bars show the standard deviation. **C.** Same as B. but for the average fraction of visited patches. **D.** Average fraction of time spent inside patches of distinct substrates (from left to right: sucrose, yeast, apple juice) for rovers (solid line, filled circles) and sitters (dashed line, open circles) as a function of the number of patches. Each point is the average of 30 larvae (30 simulation runs each). Bars show the standard deviation. **E.** Same as D. but for the average fraction of visited patches.

